# A single-channel EEG classification system for multiscale characterization of mouse vigilance state

**DOI:** 10.64898/2026.08.14.744824

**Authors:** Yazan Waddah Khaled Zaid, Pawel Matulewicz, Svenja L. Kreis, Thomas Fenzl, Anna C. Elbs, Leesa Joyce, Azra Durmic-Basic, Armin Schmuck, Mohadeseh Ragerdikashani, Sadegh Rahimi, Taro Tezuka

**Author notes:** **Correspondence:** Sadegh Rahimi, Taro Tezuka. These authors contributed equally (Sadegh Rahimi, Taro Tezuka).

## Abstract

Long-term analysis of mouse sleep is constrained by the dependence of conventional scoring on expert interpretation of electroencephalographic (EEG) and electromyographic (EMG) recordings. We developed a channel-agnostic, EEG-only framework that combines cross-animal sleep-stage classification, causal temporal organization, and probabilistic hypnodensity analysis from a single cortical EEG signal. Motor, somatosensory, and visual cortical recordings were treated independently by a convolutional–recurrent classifier, and generalization was evaluated using nested leave-one-mouse-out cross-validation in eight mice, with each test animal excluded from training, normalization, and model selection. The primary model achieved 0.897 ± 0.058 accuracy and 0.856 ± 0.076 macro-F1 across previously unseen animals while preserving the principal features of expert EEG/EMG-supported sleep architecture. Causal temporal smoothing reduced fragmented predictions and restored physiologically coherent episode durations, counts, and transition structure. Beyond categorical staging, the 4-s causal EEG window was advanced in 1-s steps to generate continuous Wake, NREM, and REM hypnodensity profiles. This representation preserved overall classification performance while revealing increased probability ambiguity and state mixing around expert-defined sleep transitions. The framework was subsequently deployed without supervised adaptation in six additional mice with 32–33 recorded days per animal, where it retained organized daily sleep architecture and probabilistic sleep structure over extended recordings while remaining sensitive to changes in recording conditions. Together, these results establish a single-channel EEG framework for robust cross-animal sleep staging, physiologically structured long-term analysis, and second-by-second characterization of sleep-state probabilities in mice.

## Introduction

Sleep staging provides the temporal framework for studying vigilance-dependent brain activity and for experiments in which observation or intervention is restricted to wakefulness, NREM sleep, or REM sleep. In mice, these states are conventionally identified by visual inspection of EEG and EMG signals. Expert scoring remains an important physiological reference, although the time required for manual annotation becomes increasingly restrictive as recordings extend from hours to days or weeks.

Automated sleep staging has substantially increased the scale at which rodent EEG can be analyzed [Sunagawa et al., 2013, Tezuka et al., 2021, Wang et al., 2023]. Deep-learning approaches have further enabled classification directly from physiological signals and incorporation of temporal information across successive epochs. Single-channel EEG is particularly attractive for long-term and previously acquired datasets because it can recover sleep information when EMG or additional cortical channels are unavailable. Previous work demonstrated accurate classification of Wake, NREM, and REM from a single mouse EEG channel and subsequently used EEG-only classification for real-time REM-specific intervention [Tezuka et al., 2021, Koyanagi et al., 2023]. High-frequency EEG activity also contains information relevant to vigilance-state discrimination, including separation of REM sleep from wakefulness [Rahimi et al., 2023]. A central remaining challenge is to establish how reliably such models generalize to completely unseen animals and whether their predictions preserve physiologically meaningful sleep organization when deployed over extended recordings.

Conventional automated scoring ultimately reduces each epoch to a single categorical stage. The underlying model probabilities contain a richer description of the evolving sleep state. Hypnodensity representations preserve the probability assigned to every sleep stage over time and have revealed graded stage transitions, sleep-stage overlap, and scoring ambiguity that are compressed in conventional hypnograms [Stephansen et al., 2018, Bakker et al., 2023]. High-frequency sleep-stage representations have similarly shown that probabilistic sleep information can be resolved at temporal scales finer than the scoring epochs used during model training [Perslev et al., 2021]. These developments provide a framework for examining sleep as a continuously evolving probabilistic process and for deriving measures of state ambiguity and state mixing in addition to conventional sleep macrostructure.

The temporal organization of the categorical hypnogram remains equally important. Isolated short predictions can have little influence on epoch-level accuracy while substantially altering episode number, duration, and transition structure. Incorporating recent temporal information into stage probabilities therefore provides a physiologically motivated means of stabilizing the predicted sequence. For long-duration recordings, the combination of robust cross-animal classification, temporal organization of predicted bouts, and probabilistic hypnodensity analysis offers complementary views of sleep macrostructure and microstructure.

In the present study, we developed a channel-agnostic, EEG-only framework for Wake, NREM, and REM classification in mice. Motor, somatosensory, and visual cortical EEG recordings were treated as independent single-channel inputs, with expert EEG/EMG-supported scoring used as the reference. Generalization was evaluated by nested leave-one-mouse-out cross-validation, keeping every test animal independent of model fitting, normalization, and model selection. We then applied a causal temporal smoothing procedure based on recent stage probabilities and examined its effect on sleep-bout organization. The same classifier was further used to generate a 1-s hypnodensity by advancing the 4-s causal EEG window in 1-s steps, providing continuous Wake, NREM, and REM probabilities from which stage ambiguity and pairwise state mixing were quantified. Finally, the framework was deployed in a separate cohort with 32–33 recorded days per animal to examine the preservation of sleep architecture and probabilistic sleep structure during long-term single-channel EEG monitoring.

## Materials and Methods

### Study cohorts

The labeled dataset initially comprised nine mice recorded continuously for approximately 23 h at a sampling frequency of 1000 Hz. One recording, M2LJ06, was excluded before model development following expert assessment of recording quality, leaving eight mice for analysis. Each recording contained EEG signals from the motor, somatosensory, and visual cortices together with EMG activity. Expert-scored epochs comprised approximately 48% wakefulness, 47% NREM sleep, and 5% REM sleep.

A separate long-term cohort comprised six mice with one EEG channel per animal. Each animal contributed 32–33 recorded days distributed over an approximately 40-day period. These recordings contained no manual sleep-stage labels and were used for long-term deployment, sleep-architecture analysis, and probabilistic hypnodensity analysis. No long-term recording contributed to supervised training, model selection, or normalization fitting.

### Expert sleep-stage scoring

The labeled recordings were divided into non-overlapping 4-s epochs. A single specialist assigned each epoch to wakefulness, NREM sleep, or REM sleep by visual inspection of the EEG and EMG signals. These expert labels served as the physiological reference for model development and evaluation. The classifier itself received EEG alone.

### Single-channel EEG organization

Each cortical EEG channel was treated as an independent single-channel example. Motor, somatosensory, and visual EEG signals were never presented simultaneously to the network, and cortical channel identity was not supplied as an input. All channels belonging to the same mouse remained within the same training, validation, or test partition.

Each 4-s epoch contained 4000 EEG samples. No additional filtering, detrending, or amplitude clipping was applied in the classifier pathway. At the 1000-Hz sampling frequency, the model therefore received signal content across the recorded spectrum up to the 500-Hz Nyquist frequency.

### Signal normalization

Standardization parameters were estimated exclusively from the training animals available within each model fit. For each within-epoch sample position *t*, the mean *µ*_*t*_ and standard deviation *σ*_*t*_ were calculated across all training channel–epoch examples. An EEG value *x*_*e,t*_, where *e* denotes one channel–epoch example, was standardized as

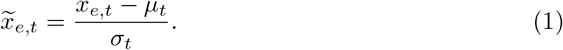

The resulting transformation was applied unchanged to validation, outer-test, and long-term recordings. Thus, normalization parameters for every held-out animal were derived entirely from other mice.

### Sleep-stage classification model

The classifier combined a one-dimensional convolutional encoder with a long short-term memory network [Hochreiter and Schmidhuber, 1997]. The encoder contained five convolutional blocks with 256, 128, 64, 32, and 16 output channels. Each block incorporated convolution, batch normalization, rectified linear activation, dropout, and temporal pooling. The resulting EEG representation was passed to a one-layer unidirectional LSTM with 128 hidden units, followed by a linear layer and three-class softmax output representing the probabilities of Wake, NREM, and REM.

The primary classifier incorporated 45 consecutive 4-s epochs, corresponding to 180 s of causal temporal context. The convolutional encoder was initialized from a model trained with a five-epoch temporal context using the same development-mouse partition. Encoder weights and batch-normalization statistics were subsequently fixed, and a new recurrent layer and classification head were trained using the longer 45-epoch sequences. The resulting model estimated the sleep stage of the most recent epoch using only present and preceding EEG information.

Training used class-weighted cross-entropy loss and the Adam optimizer with a learning rate of 10^*−*3^, weight decay of 10^*−*4^, and a batch size of 128. Dropout was set to The implementation used PyTorch [Paszke et al., 2019] and scikit-learn [Pedregosa et al., 2011].

### Nested leave-one-mouse-out evaluation

Generalization was evaluated using nested leave-one-mouse-out cross-validation. In each of eight outer folds, one mouse was retained as the final test animal and the remaining seven mice formed the development set. Full inner leave-one-mouse-out validation was performed within the development set, with six mice used for training and one for validation in each inner fold.

The inner validation procedure determined the training duration before the model was refitted using all seven development mice and evaluated once on the untouched outer-test mouse. The outer-test animal contributed no information to gradient updates, normalization fitting, class-weight calculation, checkpoint selection, or determination of training duration.

Predictions were evaluated independently for each EEG channel. For mouse-level evaluation, the class probabilities obtained from all available channels of the same animal were averaged before assignment of the final stage. This soft-voting procedure combined independently classified single-channel probability estimates while preserving the single-channel nature of the model input.

### Causal temporal smoothing and sleep-sequence correction

The network output was refined using a fixed causal smoothing procedure based on the current and two immediately preceding sleep-stage probability vectors. Let **p**_*t*_ = (*p*_*t*1_, *p*_*t*2_, *p*_*t*3_) denote the predicted probabilities of Wake, NREM, and REM at epoch *t*. The uncertainty of the current prediction was quantified using normalized Shannon entropy,

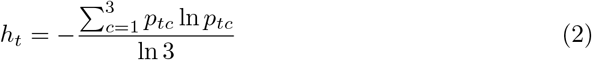

where *h*_*t*_ ranges from 0 for a maximally confident prediction to 1 when equal probability is assigned to all three stages. The contribution of preceding predictions was determined from this uncertainty and capped at 0.5,

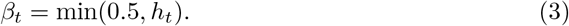

For epochs with two preceding predictions available, the temporally smoothed probability vector was calculated as

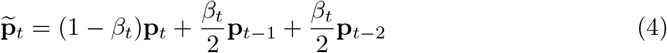

Thus, confident current predictions received little temporal adjustment, whereas predictions with greater uncertainty received increasing support from the recent stage history, with previous epochs contributing at most one half of the total weight. At the beginning of a recording, the same causal rule was applied using only the preceding probability vectors that were available. The categorical stage was then assigned as

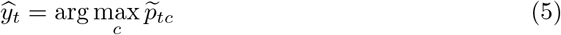

Because the procedure uses only the current and preceding predictions, no information from future epochs contributes to the corrected stage sequence.

For the labeled cohort, probability smoothing was followed by correction of REM episodes lasting no more than three consecutive epochs (12 s). These short REM segments were merged into the surrounding stage sequence, while Wake and NREM episodes were not duration-corrected. For long-term inference, the predefined minimum-duration correction was applied to Wake, NREM, and REM episodes lasting no more than 1, 3, and 4 epochs, corresponding to 4, 12, and 16 s, respectively.

Temporal correction was evaluated using episode number, episode duration, short-episode prevalence, stage-transition rates, and epoch-level classification performance.

### Hypnodensity estimation

To retain the full sleep-stage probability structure at a finer temporal sampling interval, the validated classifier was additionally evaluated using the same 4-s EEG input window advanced through each recording in 1-s steps. Thus, the input duration remained 4 s, while a new Wake, NREM, and REM probability estimate was generated every second. At each step, the 4-s causal window comprised the current 1-s interval together with the immediately preceding 3 s of EEG. This window was evaluated within the same longer causal temporal context used by the primary classifier, preserving information from preceding EEG epochs while increasing the temporal sampling of the model output.

For each second *t*, the classifier produced a three-stage probability vector

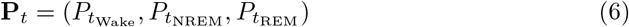

The ordered sequence of these probability vectors constituted the hypnodensity representation [Stephansen et al., 2018, Bakker et al., 2023], providing a continuous description of the relative probability assigned to each vigilance state throughout the recording.

For the labeled cohort, each mouse was processed using the model checkpoint and normalization parameters from the corresponding outer fold in which that animal had served as the held-out test mouse. The 1-s estimates therefore remained independent of model fitting and normalization for that animal. To assess whether the finer output interval altered overall classification behavior, each 1-s estimate was compared with the expert-scored 4-s epoch containing that second, and agreement was summarized using accuracy, macro-F1 score, and Cohen’s *κ*.

The same 4-s window with 1-s advancement was used for hypnodensity analysis of the long-term recordings, with model weights and normalization parameters fixed from the labeled cohort. The resulting probability sequences were subsequently used to quantify probability-weighted stage occupancy, entropy-based stage ambiguity, and pairwise state mixing.

### Hypnodensity-derived sleep microstructure

Three complementary measures were extracted from the hypnodensity. First, probability-weighted stage occupancy was calculated as the mean probability assigned to each vigilance state over the analyzed recording. This retained contributions from partially expressed stage probabilities that are removed when the probability vector is reduced to a single categorical label.

Second, stage ambiguity was quantified at each second using the Shannon entropy of the Wake, NREM, and REM probability distribution,

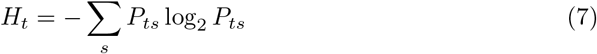

where *s* denotes the three vigilance states. Higher entropy indicates a more distributed probability assignment across states. In the labeled cohort, ambiguity was examined in relation to expert-defined sleep-stage transitions by aligning the 1-s probability sequence to transition times and comparing the temporal profile around transitions with stable portions of the recording.

Third, pairwise state mixing was quantified for Wake–NREM, NREM–REM, and Wake–REM using the probability overlap between each pair of stages,

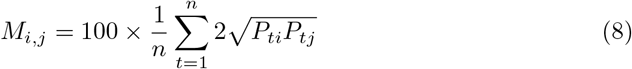

where *i* and *j* denote a pair of vigilance states and *n* is the number of 1-s probability estimates. Larger values indicate greater simultaneous probability assigned to the two states across the analyzed recording.

### Prediction-based channel screening

The primary classification analysis included all available EEG channels. A secondary prediction-based screen was examined to identify channels associated with biologically implausible full-recording stage distributions. A channel was flagged when the predicted NREM fraction was below 0.20 or the predicted REM fraction was below 0.02.

Screening was performed at the channel level, used no expert labels, and was defined after inspection of the model outputs. The unscreened analysis remained the primary estimate of cross-animal performance.

### Classification and sleep-architecture measures

Classification performance was evaluated using accuracy, macro-averaged F1 score, Cohen’s *κ*, and stage-specific sensitivity, specificity, precision, and F1 score. Confusion matrices were pooled across the predictions obtained when each mouse served as the outer-test animal.

Sleep architecture was evaluated from the complete categorical stage sequence. For each state, total duration, episode number, mean episode duration, median episode duration, and the proportion of episodes lasting no more than 12 s were calculated.

Stage-transition rates were expressed as transitions per hour.

### Long-term sleep-architecture and hypnodensity analysis

The long-term cohort was analyzed using the 45-epoch deployment model with fixed batch-normalization statistics derived from the labeled data. Model weights and normalization parameters were held constant throughout long-term inference.

Daily Wake, NREM, and REM proportions were calculated separately for every recorded day. Episode number was expressed per 24 h, and mean and median episode durations were calculated for each state. Measurements from the expert-scored cohort were displayed as a physiological reference for interpretation of the long-term predictions.

The recording system changed between recording days 15 and 17, with calendar day 16 absent from the recordings. Long-term sleep architecture was therefore summarized separately before and after this transition. Missing calendar days remained missing and were not interpolated.

Hypnodensity microstructure was examined on one representative pre-change day from each long-term animal. The representative day was selected by a fixed rule as the pre-change recording day whose predicted stage composition was closest to that animal’s median pre-change composition. Each selected 24-h recording was processed at 1-s cadence using the same causal sliding-window procedure applied to the labeled cohort. Probability-weighted stage occupancy, mean ambiguity, and pairwise state-mixing indices were then calculated for each animal.

### Statistical analysis

The mouse was treated as the independent biological unit throughout the analysis.Classification performance was summarized across the eight outer-test mice as mean ± standard deviation. Expert- and model-derived measures of sleep architecture were compared within animals using paired Wilcoxon signed-rank tests, with *p <* 0.05 considered statistically significant.

Hypnodensity measures were first calculated independently for each outer-test mouse and then summarized across the cohort. Transition-related changes in entropy and stage disagreement were quantified according to temporal distance from expert-defined stage transitions, including the predefined ± 2-s transition interval and stable periods.

Long-term sleep architecture and hypnodensity microstructure were summarized at the animal level using descriptive statistics, reflecting the absence of expert-scored labels in the long-term cohort.

## Results

### Cross-animal validation of EEG-only sleep-stage classification

Cross-animal performance was evaluated in eight labeled mice (LM), with each animal serving once as an independent outer-test subject. The corrected stage sequences preserved the major temporal organization of expert EEG/EMG-supported scoring across complete recordings. Total Wake, NREM, and REM duration showed similar cohort-level distributions between expert and corrected scoring. Episode counts and mean episode durations showed greater animal-to-animal dispersion in the corrected sequences. Paired Wilcoxon signed-rank tests identified no significant differences in total stage duration, episode count, or mean episode duration across the eight animals. LM M2LJ05, selected by the predefined median macro-F1 rule, provides a representative example of the correspondence between expert and corrected sleep organization (Figure 1).

**Figure 1.**
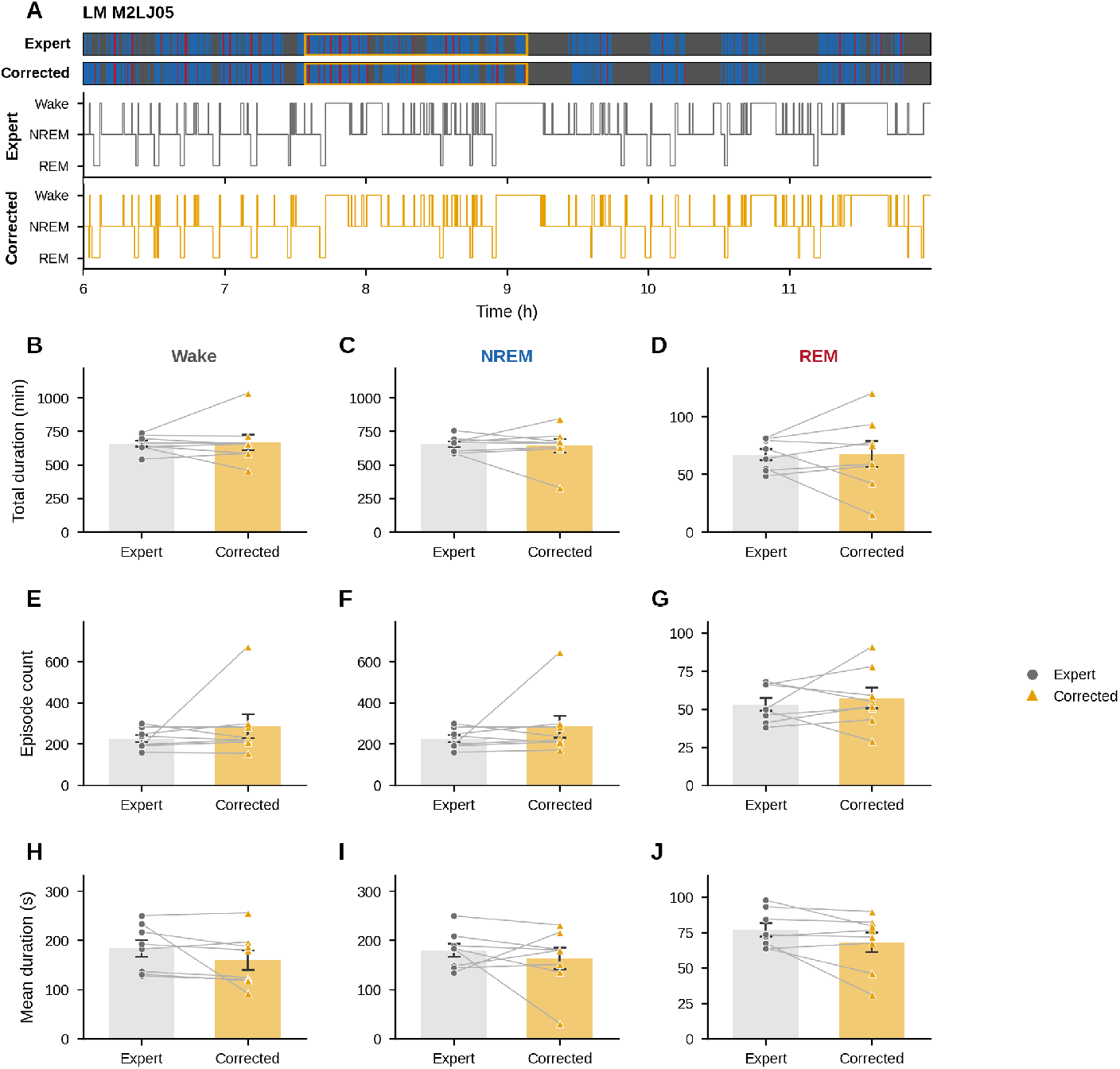
Expert and corrected sleep architecture in the labeled cohort. (A) Expert and corrected hypnograms for LM M2LJ05. The upper tracks represent the complete recording and the outlined interval identifies the 6-h segment enlarged below. (B–D) Total duration of Wake, NREM, and REM. (E–G) Number of episodes per recording. (H–J) Mean episode duration. Circles indicate Expert values and triangles indicate Corrected values; lines connect measurements from the same animal. Bars and error bars represent mean ± SEM across eight labeled mice. LM denotes labeled mouse.

Across the eight outer-test mice, the primary unscreened model achieved an accuracy of 0.897 ± 0.058, macro-F1 of 0.856 ± 0.076, and Cohen’s *κ* of 0.808 ± 0.115. Wake and NREM showed mean F1 scores of 0.904 ± 0.054 and 0.897 ± 0.077, respectively. REM performance showed greater inter-animal variation, with an F1 score of 0.767 ± 0.119, sensitivity of 0.806 0.206, and specificity of 0.987 ± 0.010. Pooled predictions from 165,180 previously unseen 4-s epochs yielded sensitivities of 91.0% for Wake, 89.2% for NREM, and 81.7% for REM. Most classification errors involved Wake and NREM, and direct Wake–REM errors were less frequent (Figure 2).

**Figure 2.**
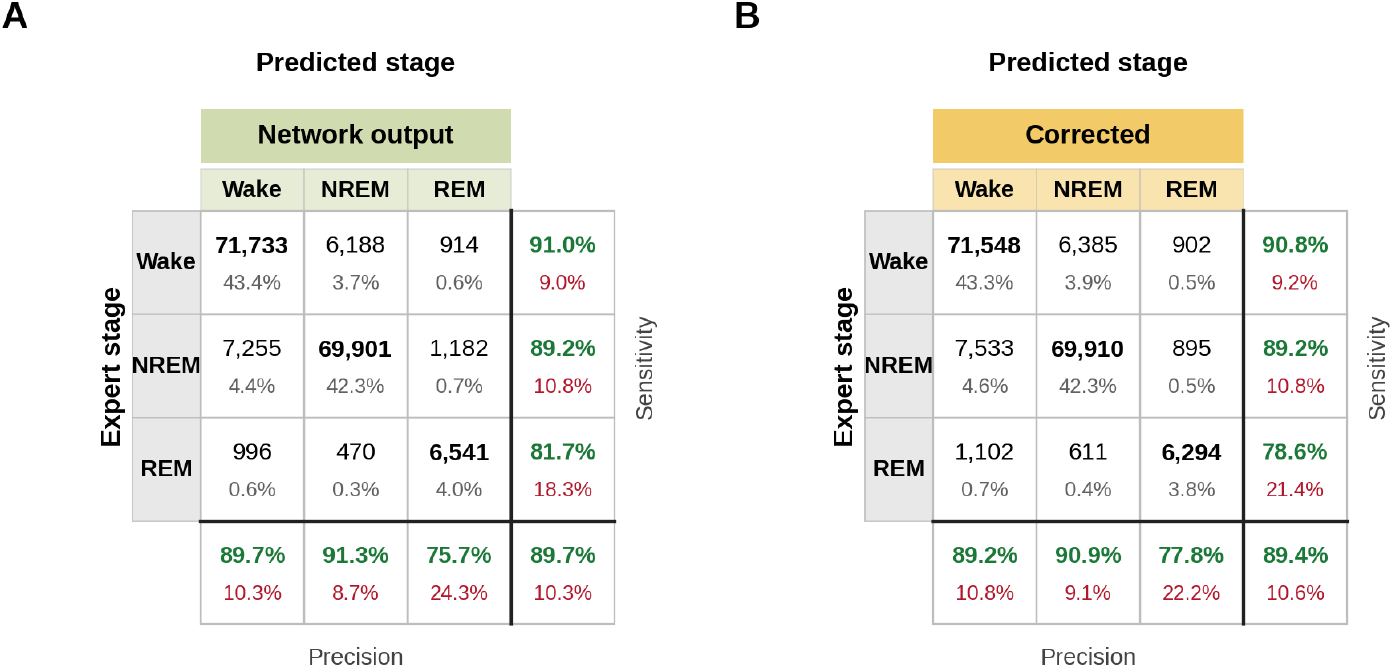
Nested cross-animal sleep-stage classification performance. Pooled confusion matrices from the eight outer-test mice comprising 165,180 previously unseen 4-s epochs. (A) Network output. (B) Corrected output following causal temporal smoothing and sleep-sequence correction. Rows correspond to Expert stages and columns to predicted stages. Each matrix cell contains the epoch count and its percentage of the complete pooled dataset. Green and red values on the right indicate stage sensitivity and the corresponding error fraction. Green and red values below the matrix indicate precision and the corresponding error fraction for each predicted stage.

Temporal correction produced small changes in the aggregate epoch-level performance measures (Table 1). A secondary prediction-based channel screen increased accuracy to 0.917 ± 0.037, macro-F1 to 0.892 ± 0.034, REM F1 to 0.835 ± 0.041, and REM sensitivity to 0.902 ± 0.051. The unscreened evaluation remained the primary analysis because the screening thresholds were defined after examination of the model outputs.

**Table 1.** Sleep-stage classification across eight outer-test mice.

| Measure | Network output | Corrected |
| --- | --- | --- |
| Accuracy | $0.897 \pm 0.058$ | $0.895 \pm 0.060$ |
| Macro-F1 | $0.856 \pm 0.076$ | $0.849 \pm 0.091$ |
| Cohen's $\kappa$ | $0.808 \pm 0.115$ | $0.802 \pm 0.121$ |
| Wake F1 | $0.904 \pm 0.054$ | $0.901 \pm 0.055$ |
| NREM F1 | $0.897 \pm 0.077$ | $0.894 \pm 0.081$ |
| REM F1 | $0.767 \pm 0.119$ | $0.752 \pm 0.160$ |
| REM sensitivity | $0.806 \pm 0.206$ | $0.773 \pm 0.242$ |
| REM specificity | $0.987 \pm 0.010$ | $0.989 \pm 0.009$ |

Values are mean ± SD across the eight outer-test mice. The primary analysis includes all available EEG channels before prediction-based channel screening.

### Temporal correction and sleep-episode architecture

The direct network output produced frequent short stage assignments that fragmented the predicted sleep sequence. Temporal correction reduced episode counts across all three stages and increased mean episode duration toward the distributions obtained from expert scoring. The change was most pronounced for REM, where correction substantially reduced the excess of short episodes. Wake and NREM showed corresponding reductions in short-event fragmentation. The principal transition pathways remained Wake–NREM and NREM–REM after correction (Figure 3).

**Figure 3.**
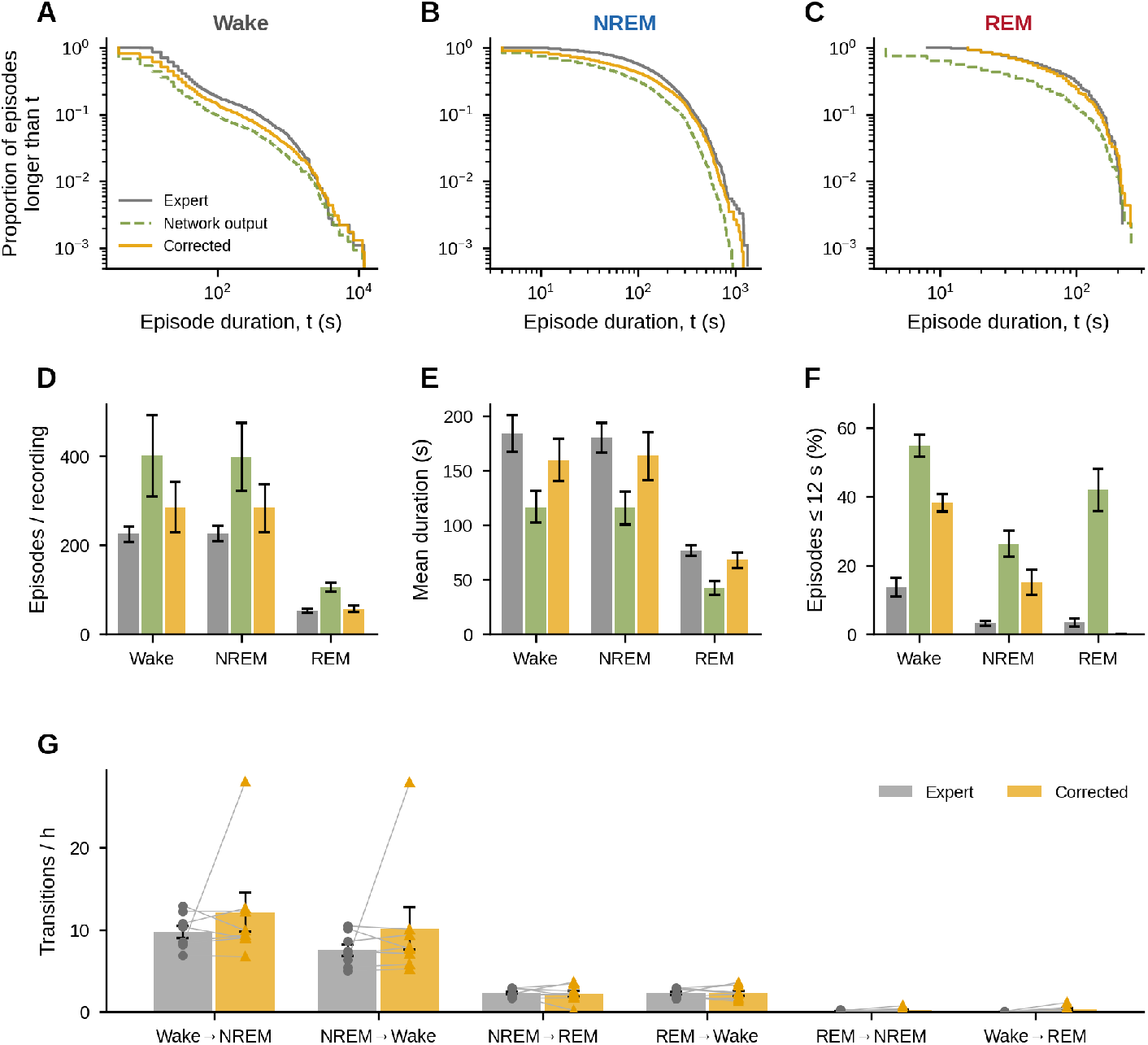
Effect of temporal correction on sleep-episode architecture. (A–C) Proportion of Wake, NREM, and REM episodes with duration greater than (t) for Expert scoring, Network output, and Corrected sequences. (D) Number of episodes per recording. (E) Mean episode duration. (F) Percentage of episodes lasting no more than 12 s. Bars and error bars in D–F represent mean ± SEM across eight labeled mice. (G) Stage-transition rates for Expert and Corrected sequences, expressed as transitions per hour; individual animals are shown together with the group mean ± SEM.

The effect on sleep architecture was considerably larger than the effect on epoch-level classification. This relationship indicates that isolated classification changes can alter episode number, duration, and transition structure without materially changing overall accuracy.

### Hypnodensity analysis of transition-associated uncertainty

The complete Wake, NREM, and REM probability output was retained to examine sleep-state dynamics at a finer temporal sampling interval. The same 4-s causal EEG window was advanced in 1-s steps, producing one three-stage probability estimate per second. Across the eight outer-test mice, the resulting 1-s categorical sequence achieved a mean accuracy of 0.893, macro-F1 of 0.850, and Cohen’s *κ* of 0.800. These values closely followed the corresponding native 4-s classification results.

Stable periods were characterized by sustained dominance of one stage probability. Changes in expert-defined stage were accompanied by redistribution of probability among Wake, NREM, and REM and increased entropy of the three-stage probability vector. The full-recording and enlarged probability trajectories from LM M2LJ05 illustrate this structure (Figure 4).

**Figure 4.**
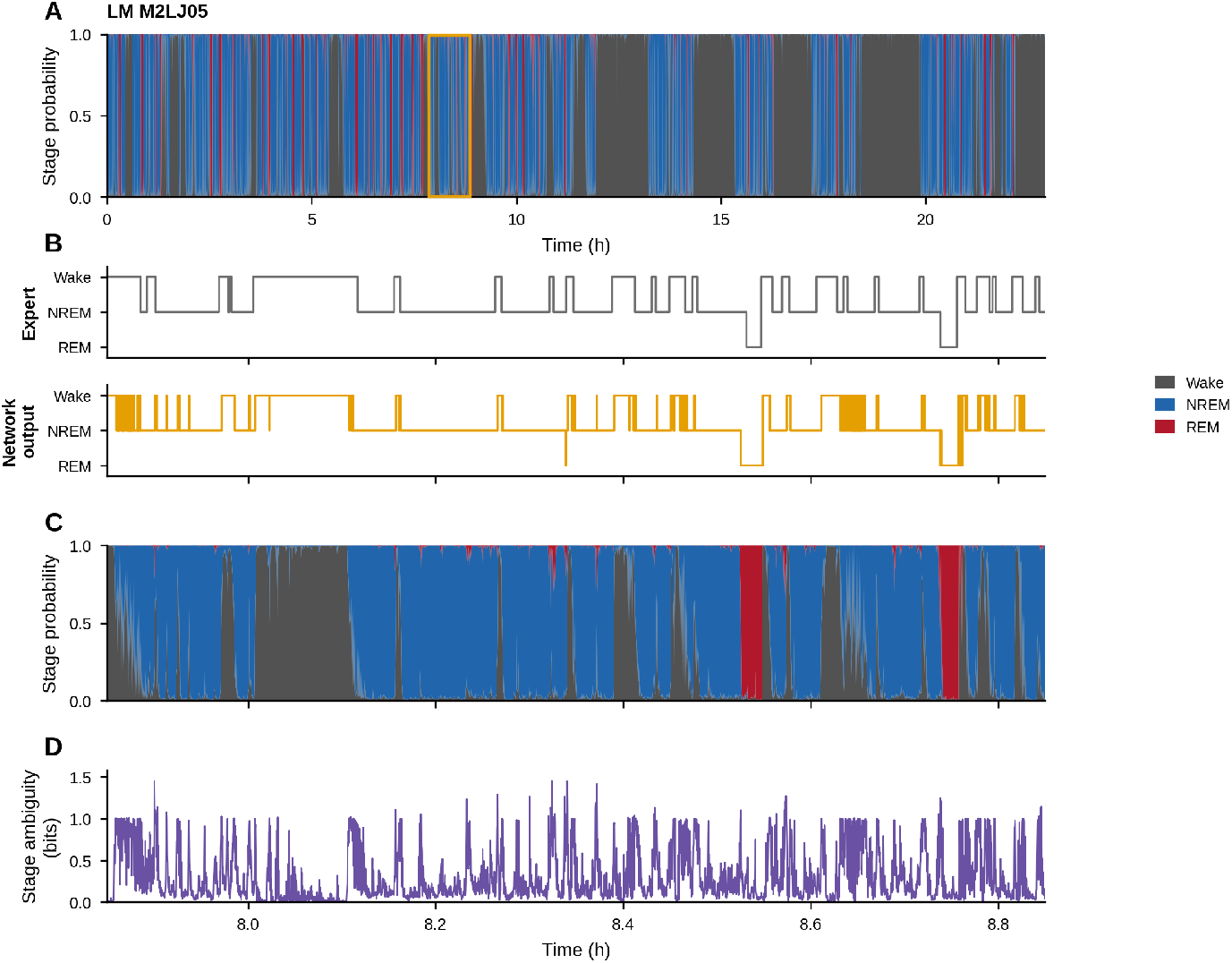
Hypnodensity analysis in the labeled cohort. (A) Wake, NREM, and REM probability trajectories across the complete recording from LM M2LJ05; the outlined region identifies the interval enlarged below. (B) Expert 4-s sleep-stage sequence and Network output generated at 1-s intervals from the same 4-s causal EEG window advanced in 1-s steps. (C) Wake, NREM, and REM probabilities over the selected interval. (D) Shannon entropy of the three-stage probability vector over the same interval, expressed in bits.

Transition-aligned analysis showed a reproducible temporal increase in model uncertainty around expert-defined stage boundaries. Mean stage ambiguity increased through the transition and reached its maximum several seconds after the annotated boundary. Disagreement between the 1-s network assignment and the corresponding expert-scored 4-s epoch increased sharply at the transition and declined over the following seconds. The temporal pattern was retained across individual mice and in the cohort mean (Figure 5). Mean hypnodensity ambiguity across the complete labeled recordings was 0.307 bits. Pairwise probability overlap was greatest for Wake–NREM, followed by NREM–REM and Wake–REM.

**Figure 5.**
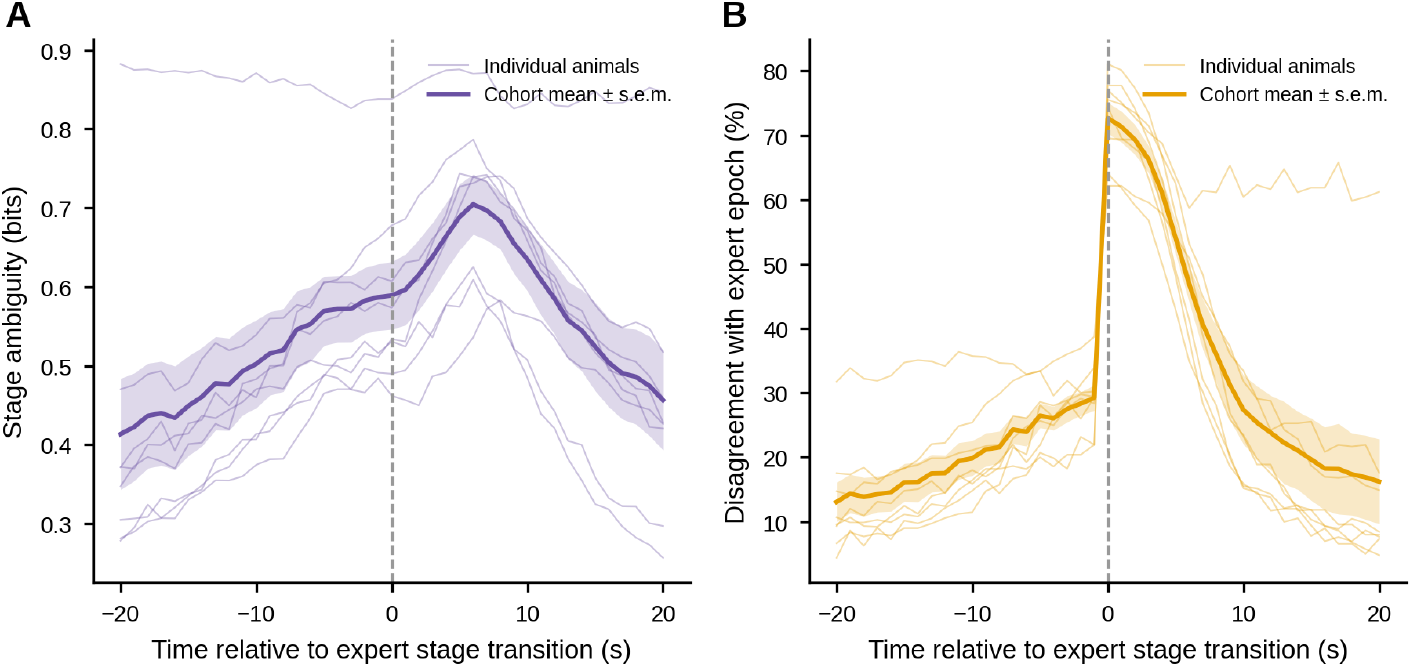
Sleep-stage uncertainty around expert-defined transitions. (A) Shannon entropy of the Wake, NREM, and REM probability vector as a function of time relative to expert-defined stage transitions. (B) Percentage of 1-s Network output assignments that differed from the expert-scored 4-s epoch containing the corresponding second. Time zero denotes the expert-defined stage transition. Thin lines represent individual labeled mice. Thick lines represent the cohort mean and shaded regions indicate mean ± SEM across eight animals.

### Long-term sleep architecture and recording-system sensitivity

The classifier was deployed without supervised adaptation in six long-term mice (LTM), each contributing 32–33 recorded days distributed across an approximately 40-day period. Sleep episodes remained organized across repeated recording days and showed stable animal-specific patterns over extended intervals. Mean REM episode duration frequently occupied the range observed in the expert-scored cohort. NREM episode counts also commonly overlapped the labeled reference distribution. Wake and NREM episode durations showed broader between-animal variation, and REM episode counts varied substantially across the long-term cohort (Figure 6).

**Figure 6.**
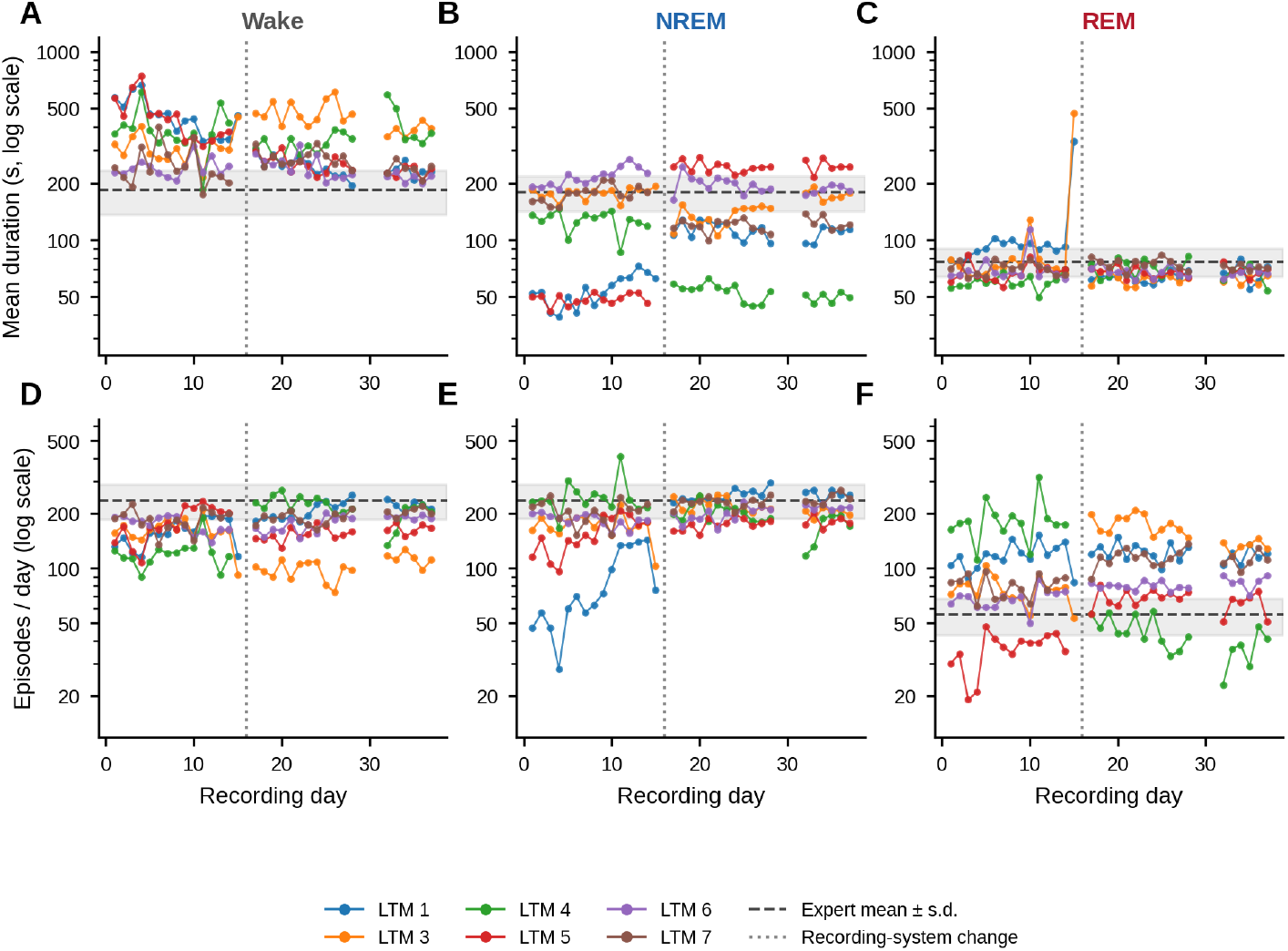
Sleep-episode architecture across long-term recordings. (A–C) Daily mean episode duration for Wake, NREM, and REM in six long-term mice. (D–F) Number of episodes per recorded day for the corresponding stages. Y-axes are logarithmically scaled. Each colored line represents one long-term mouse. Dashed horizontal lines indicate the labeled-cohort mean and grey bands indicate mean ± SD. The vertical dotted line denotes the recording-system change between recording days 15 and 17. Lines are interrupted on calendar days without recordings. LTM denotes long-term mouse.

The recording-system change coincided with marked redistribution of predicted Wake and NREM in several animals. LTM 1 showed a large decrease in Wake and corresponding increase in NREM after the change. LTM 4 showed a large increase in Wake and reduction in NREM. LTM 5 also showed a pronounced redistribution of Wake and NREM. LTM 6 showed comparatively stable stage composition across the two recording blocks. REM changes were smaller in absolute daily proportion for most animals. The direction and magnitude of the recording-associated changes were animal specific (Figure 7).

**Figure 7.**
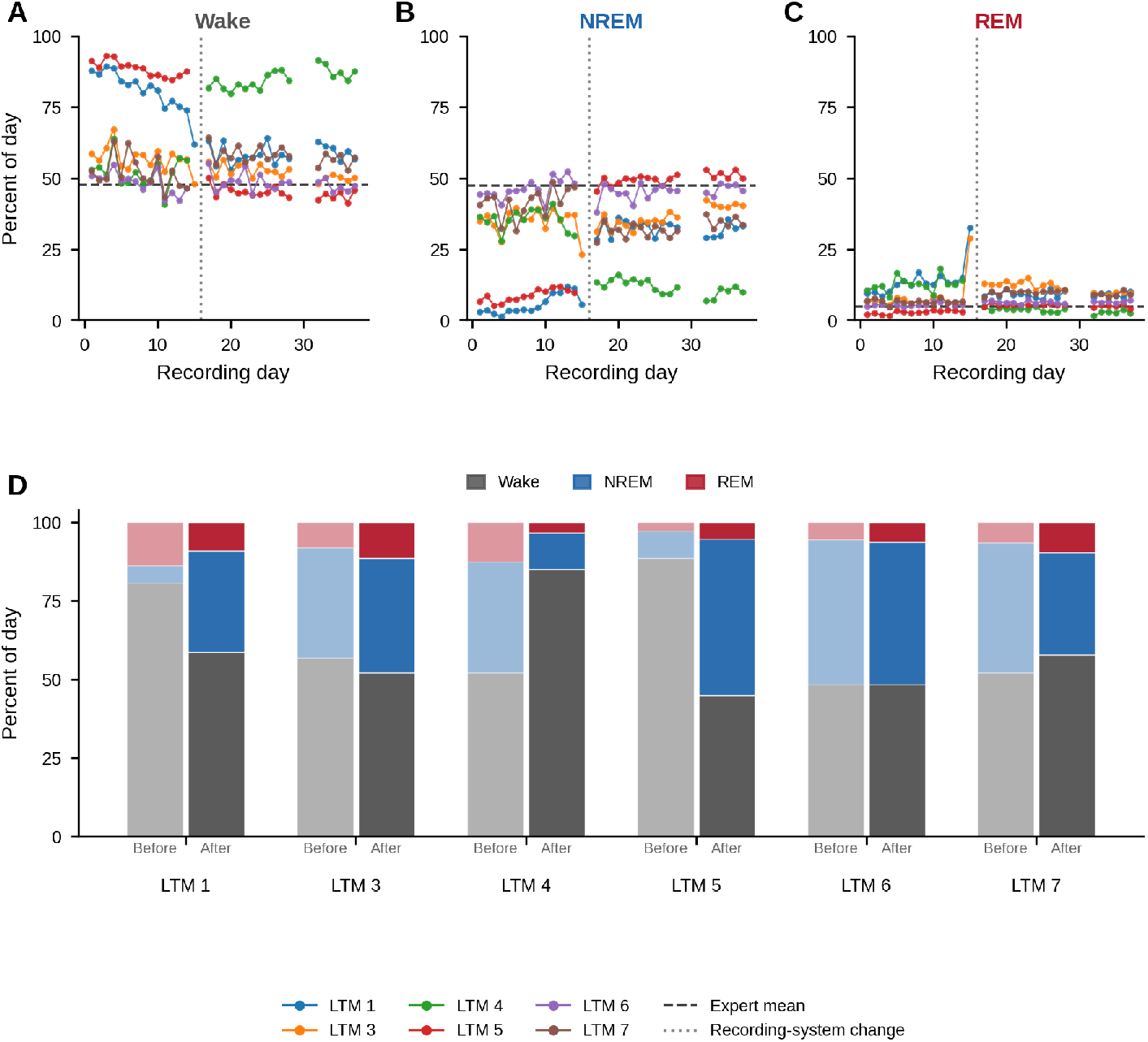
Sleep-stage composition across the recording-system change. (A–C) Daily percentage of Wake, NREM, and REM across the long-term recording period for each of the six animals. Dashed horizontal lines indicate the corresponding labeled-cohort mean. The vertical dotted line denotes the recording-system change between days 15 and 17, and trajectories are interrupted on calendar days without recordings. (D) Mean Wake, NREM, and REM composition for each animal before and after the recording-system change. Each pair of stacked bars represents one long-term mouse.

### Long-term hypnodensity and probabilistic sleep microstructure

Hypnodensity analysis was extended to representative pre-change recordings from the long-term cohort. The selected recording from LTM 6 retained sustained Wake, NREM, and REM probability structure across the light–dark cycle. The enlarged interval showed repeated periods of dominant stage probability together with graded probability redistribution and changes in entropy (Figure 8).

**Figure 8.**
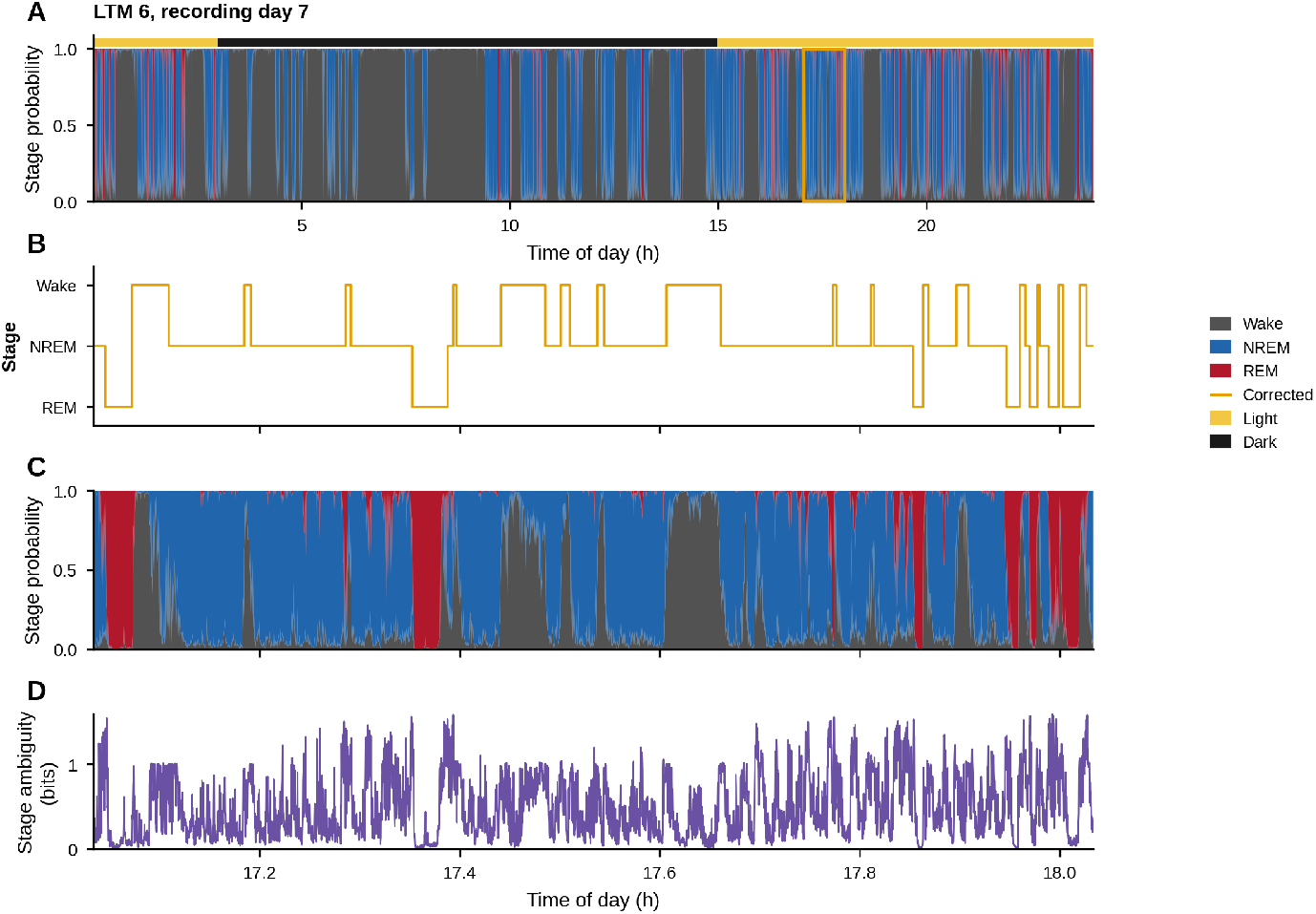
Hypnodensity during long-term single-channel EEG recording. (A) Wake, NREM, and REM probability trajectories across recording day 7 from LTM 6, selected using the predefined representative-day rule. The light–dark bar indicates the recording phase and the outlined region identifies the interval enlarged below. (B) Corrected categorical sleep-stage sequence for the selected interval. (C) Corresponding Wake, NREM, and REM probability trajectories. (D) Shannon entropy of the three-stage probability vector over the same interval, expressed in bits.

Across representative long-term days, mean stage ambiguity was 0.365 bits, compared with 0.307 bits in the labeled cohort. Wake–NREM state mixing remained similar between cohorts, with mean indices of 34.3 in the long-term recordings and 33.2 in the labeled recordings. NREM–REM mixing averaged 9.4 in the long-term cohort and 7.1 in the labeled cohort. Wake–REM mixing averaged 9.9 and 5.5, respectively. The long-term animals showed greater dispersion in probability overlap involving REM, with LTM 4 contributing the highest ambiguity and the largest NREM–REM overlap (Figure 9).

**Figure 9.**
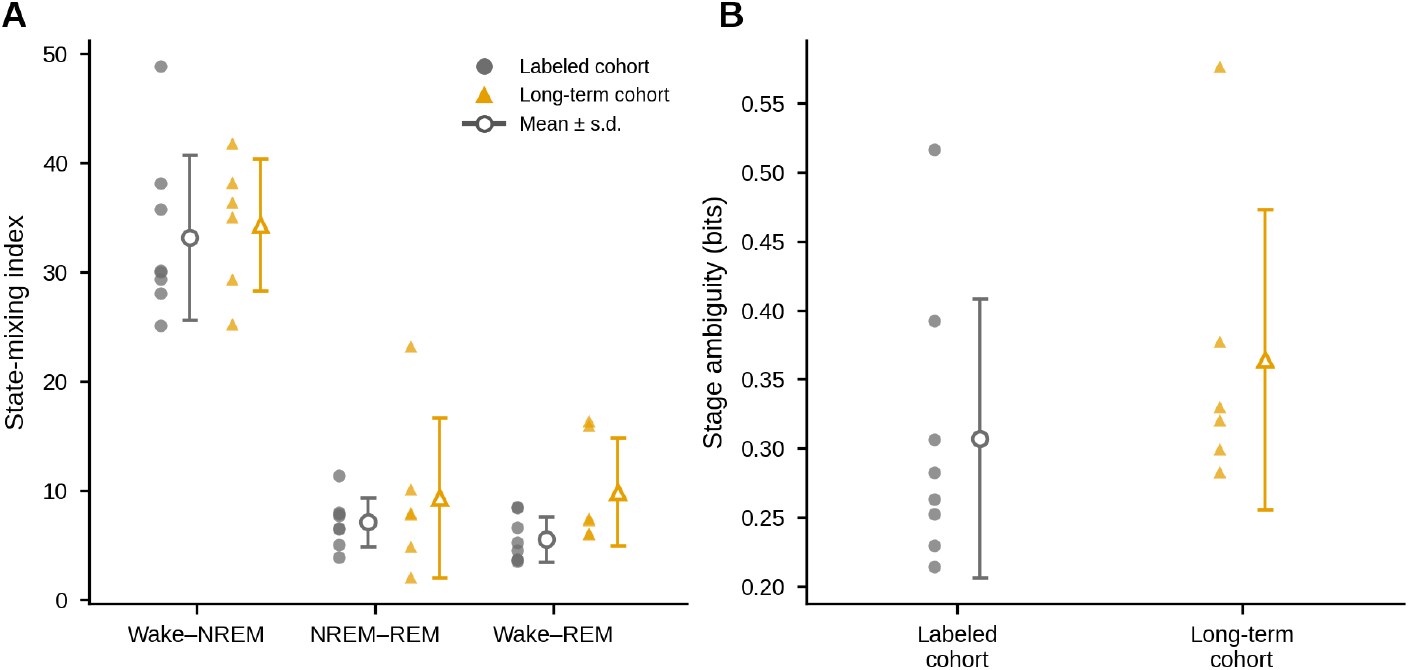
Hypnodensity-derived sleep microstructure in labeled and long-term recordings. (A) Animal-level pairwise state-mixing indices for Wake–NREM, NREM–REM, and Wake–REM in the labeled cohort and representative pre-change recordings from the long-term cohort. (B) Mean stage ambiguity for the same animals, quantified as Shannon entropy of the Wake, NREM, and REM probability vector. Filled symbols represent individual animals. Open symbols and error bars indicate cohort mean ± SD.

## Discussion

The present study developed a single-channel EEG framework for automated mouse sleep analysis across unseen animals and extended recordings. Nested leave-one-mouse-out evaluation showed stable cross-animal classification of Wake, NREM, and REM, and the resulting stage sequences preserved major features of expert-derived sleep architecture. Temporal correction improved episode organization, and hypnodensity analysis exposed the underlying stage-probability structure at 1-s output intervals. Deployment in a separate long-term cohort further demonstrated sustained sleep architecture and probabilistic organization across 32–33 recorded days per animal.

The findings extend previous single-channel mouse sleep-staging and EEG-driven intervention studies [Tezuka et al., 2021, Koyanagi et al., 2023] by combining cross-animal validation, channel-agnostic processing across multiple cortical recording sites, and long-duration deployment. Each outer-test animal remained excluded from training, normalization, and model selection, providing a stringent measure of transfer to previously unseen mice. This design is directly relevant to archived and prospective datasets in which only one cortical EEG signal is available. The classifier itself operated on one EEG channel at a time, and mouse-level evaluation in the labeled cohort combined probabilities across independently classified channels when multiple recordings were available.

REM remained the most variable stage across animals. Its low prevalence and electrophysiological similarity to wakefulness increase the difficulty of EEG-only discrimination, particularly in the absence of muscle-tone information during inference. High REM specificity and sensitivity above 0.80 in the primary analysis indicate that a single cortical EEG signal nevertheless contained substantial information for REM identification. High-frequency EEG activity may contribute to this separation, consistent with previous observations of REM–Wake discrimination at frequencies extending beyond conventional sleep bands [Rahimi et al., 2023]. The remaining inter-animal variability is particularly relevant for applications centered on REM-specific physiology or intervention.

Temporal correction and hypnodensity characterized complementary levels of sleep organization. Temporal correction reduced short fragmented assignments and shifted episode counts, durations, and transition structure toward expert-scored distributions, indicating that epoch-level classification metrics alone do not capture the quality of reconstructed sleep architecture. Hypnodensity retained the full Wake, NREM, and REM probability distribution and revealed graded probability changes around expert-defined stage boundaries. The 4-s causal EEG window was advanced in 1-s steps, preserving the native input duration while increasing the temporal sampling of model output. Entropy and stage disagreement increased around annotated transitions, with a reproducible temporal asymmetry after the boundary. Part of this transition-associated uncertainty is expected from the overlap of adjacent-stage EEG within the 4-s causal window. The resulting probability structure is consistent with the use of hypnodensity for quantifying sleep-stage ambiguity and probabilistic overlap [Stephansen et al., 2018, Bakker et al., 2023].

Long-term deployment showed persistent animal-specific sleep organization across repeated recording days and also exposed sensitivity to recording conditions. Several episode measures remained within or near the range observed in the labeled cohort, and Wake–NREM state mixing remained similar between labeled and representative long-term recordings. The recording-system change coincided with substantial redistribution of predicted Wake and NREM in several animals, with different directions and magnitudes across the cohort. This observation supports routine inspection of signal characteristics, stage composition, episode structure, and prediction confidence during extended deployment. Prediction-based channel screening produced improved classification in the labeled cohort and may provide an additional quality-control layer after independent validation of its thresholds.

Several limitations define the present scope. The labeled cohort contained eight analyzed mice and one expert scorer, precluding assessment of inter-rater variability. Baseline mouse-level summaries combined independently classified channels when several cortical recordings were available, whereas each long-term animal contributed a single EEG channel. The long-term cohort lacked expert stage labels, so its evaluation was descriptive and based on sleep architecture, temporal organization, and probability structure. Hypnodensity in the long-term cohort was examined on representative pre-change days rather than continuously across all recorded days. The recording-system change also limits biological interpretation of changes across that boundary. Finally, the present evaluation involved healthy mice, and pathological or pharmacologically altered EEG may require dedicated validation.

Single-channel EEG analysis can increase the value of existing recordings in which EMG or additional EEG channels were not collected. Reuse of such datasets may reduce the need to repeat animal experiments solely to obtain additional physiological signals, contributing to the Reduction principle of the 3Rs framework [Russell and Burch, 1959]. Further work should incorporate targeted expert scoring of long-term periods, validation across recording conditions, independent assessment of channel-quality criteria, and evaluation in pathological EEG datasets. These steps will determine the broader applicability of the framework for long-term sleep analysis in experimental models of neurological disease.

## Supporting Information

Supporting information will be provided as separate files accompanying the manuscript.

### S1 Data

#### Source data for the main figures

This file contains the numerical values underlying the main figures, including per-animal classification results, confusion matrices, sleep-episode measures, daily long-term sleep architecture, comparisons with the manually scored cohort, and hypnodensity-derived probability, ambiguity, transition, and state-mixing measures.

### S2 Data

#### Per-animal long-term sleep analysis

This file contains daily Wake, NREM, and REM proportions, episode counts and durations, light–dark summaries, representative 24-h recordings, fixed-rule 6-h excerpts, and the corresponding hypnodensity representations for individual long-term animals.

### S3 Data

#### Additional model and sensitivity analyses

This file contains per-channel classification performance, prediction-based channel screening, the five-epoch context comparison, alternative normalization conditions during long-term inference, and detailed per-animal hypnodensity and transition-related analyses.

## Acknowledgments

We thank the members of the Neuroinformatics Laboratory at the University of Tsukuba and the Institute of Pharmacology at the Medical University of Innsbruck for discussions concerning mouse sleep-stage interpretation, long-term EEG analysis, and physiological assessment of sleep architecture.

## Notes

### Competing Interest Statement

The authors have declared no competing interest.

### Funding

- Japanese Government (MEXT) Scholarship

## References

Jessie P. Bakker, Marco Ross, Andreas Cerny, Ray Vasko, Edmund Shaw, Samuel Kuna, Ulysses J. Magalang, Naresh M. Punjabi, and Peter Anderer. Scoring sleep with artificial intelligence enables quantification of sleep stage ambiguity: hypnodensity based on multiple expert scorers and auto-scoring. Sleep, 46(2):zsac154, 2023. doi: 10.1093/sleep/zsac154. URL https://doi.org/10.1093/sleep/zsac154.

Sepp Hochreiter and Jürgen Schmidhuber. Long short-term memory. Neural Computation, 9(8):1735–1780, 1997. doi: 10.1162/neco.1997.9.8.1735. URL https://doi.org/10.1162/neco.1997.9.8.1735.

Iyo Koyanagi, Taro Tezuka, Jiahui Yu, Sakthivel Srinivasan, Toshie Naoi, Shinnosuke Yasugaki, Ayaka Nakai, Shimpei Taniguchi, Yu Hayashi, Yasushi Nakano, and Masanori Sakaguchi. Fully automatic rem sleep stage-specific intervention systems using single eeg in mice. Neuroscience Research, 186:51–58, 2023. doi: 10.1016/j.neures.2022.10.001. URL https://doi.org/10.1016/j.neures.2022.10.001.

Adam Paszke, Sam Gross, Francisco Massa, Adam Lerer, James Bradbury, Gregory Chanan, Trevor Killeen, Zeming Lin, Natalia Gimelshein, Luca Antiga, Alban Desmaison, Andreas Köpf, Edward Yang, Zachary DeVito, Martin Raison, Alykhan Tejani, Sasank Chilamkurthy, Benoit Steiner, Lu Fang, Junjie Bai, and Soumith Chintala. Pytorch: An imperative style, high-performance deep learning library. In Advances in Neural Information Processing Systems,volume 32, pages 8024–8035, 2019.

Fabian Pedregosa, Gaël Varoquaux, Alexandre Gramfort, Vincent Michel, Bertrand Thirion, Olivier Grisel, Mathieu Blondel, Peter Prettenhofer, Ron Weiss, Vincent Dubourg, Jake Vanderplas, Alexandre Passos, David Cournapeau, Matthieu Brucher, Matthieu Perrot, and Édouard Duchesnay. Scikit-learn: Machine learning in python. Journal of Machine Learning Research, 12:2825–2830, 2011. URL https://jmlr.org/papers/v12/pedregosa11a.html.

Mathias Perslev, Sune Darkner, Lykke Kempfner, Miki Nikolic, Poul Jørgen Jennum, and Christian Igel. U-sleep: resilient high-frequency sleep staging. npj Digital Medicine, 4(1):72, 2021. doi: 10.1038/s41746-021-00440-5. URL https://doi.org/10.1038/s41746-021-00440-5.

Sadegh Rahimi, Amir Soleymankhani, Leesa Joyce, Pawel Matulewicz, Matthias Kreuzer, Thomas Fenzl, and Meinrad Drexel. Discriminating rapid eye movement sleep from wakefulness by analyzing high frequencies from single-channel eeg recordings in mice. Scientific Reports, 13(1):9608, 2023. doi: 10.1038/s41598-023-36520-7. URL https://doi.org/10.1038/s41598-023-36520-7.

William Moy Stratton Russell and Rex Leonard Burch. The Principles of Humane Experimental Technique. Methuen, London, 1959.

Jens B. Stephansen, Alexander N. Olesen, Mads Olsen, Aditya Ambati, Eileen B. Leary, Hyatt E. Moore, Oscar Carrillo, Ling Lin, Fang Han, Han Yan, Yun L. Sun, Yves Dauvilliers, Sabine Scholz, Lucie Barateau, Birgit Högl, Ambra Stefani, Seung Chul Hong, Tae Won Kim, Fabio Pizza, Giuseppe Plazzi, Stefano Vandi, Elena Antelmi, Dimitri Perrin, Samuel T. Kuna, Paula K. Schweitzer, Clete Kushida, Paul E. Peppard, Helge B. D. Sørensen, Poul Jennum, and Emmanuel Mignot. Neural network analysis of sleep stages enables efficient diagnosis of narcolepsy. Nature Communications, 9:5229, 2018. doi: 10.1038/s41467-018-07229-3. URL https://doi.org/10.1038/s41467-018-07229-3.

Genshiro A. Sunagawa, Hiroyoshi Séi, Shigeki Shimba, Yoshihiro Urade, and Hiroki R. Ueda. Faster: an unsupervised fully automated sleep staging method for mice. Genes to Cells, 18(6):502–518, 2013. doi: 10.1111/gtc.12053. URL https://doi.org/10.1111/gtc.12053.

Taro Tezuka, Deependra Kumar, Sima Singh, Iyo Koyanagi, Toshie Naoi, and Masanori Sakaguchi. Real-time, automatic, open-source sleep stage classification system using single eeg for mice. Scientific Reports, 11(1):11151, 2021. doi: 10.1038/s41598-021-90332-1. URL https://doi.org/10.1038/s41598-021-90332-1.

Lei A. Wang, Ryan Kern, Eunah Yu, Soonwook Choi, and Jen Q. Pan. Intellisleepscorer, a software package with a graphic user interface for automated sleep stage scoring in mice based on a light gradient boosting machine algorithm. Scientific Reports, 13:4275, 2023. doi: 10.1038/s41598-023-31288-2. URL https://doi.org/10.1038/s41598-023-31288-2.

